# Allometric spreading and focal adhesion collective organization are coordinated by cell-scale geometrical constraints

**DOI:** 10.1101/2025.01.17.632363

**Authors:** Célian Bimbard, Ali-Alhadi Wahhod, Démosthène Mitrossilis, Joseph Vermeil, Rémi Bousquet, Alain Richert, David Pereira, Pauline Durand, Sophie Asnacios, Jocelyn Étienne, Atef Asnacios, Jonathan Fouchard

## Abstract

Focal adhesions are protein complexes that transmit actin cytoskeleton forces to the extracellular matrix and serve as signaling hubs that regulate cell physiology. While their growth is achieved through a local force-dependent process, the requirement of sustaining stress at the cell scale suggests a global regulation of the collective organization of focal adhesions. To investigate evidence of such large-scale regulation, we compared changes in cell shape and the organization of focal adhesion-like structures during the early spreading of fibroblasts either on a two-dimensional substrate or confined between two parallel plates, and for cells of different volumes. In this way, we reveal that the areal density of focal adhesions is conserved regardless of cell size or third-dimensional confinement, despite different absolute values of the surface covered by adhesion clusters. In particular, the width of the focal adhesions ring, which fills the flat lamella at the cell front, adapts to cell size and third-dimensional confinement and scales with cell-substrate contact radius. We find that this contact radius also adapts in the parallel-plate geometry so that the cumulated area of cell-substrate contact is conserved at the cell scale. We suggest that this behavior is the result of 3D cell shape changes which govern spreading transitions. Indeed, because of volume conservation constraints, the evolution of cell-body contact angle adjusts according to cell size and confinement, whereas the rate of early spreading at the cell-substrate contact is not affected by third-dimensional geometry. Overall, our data suggest that a coordination between global and local scales mediates the adaptation of cell-substrate contacts and focal adhesions distribution to large scale geometrical constraints, which allows an invariant cell-substrate adhesive energy.

## Introduction

In vivo, adherent cells encounter complex geometrical environments since the extra-cellular matrix (ECM) varies in density and topography in different organs (1, 2) and tissue physiological state (3, 4). The reconstitution of three-dimensional ECM networks *in vitro* has revealed that cell shape is influenced by the architecture of its micro-environment (4, 5). Because cell spreading modulates cellular functions such as traction force (6), differentiation (7) and cell cycle processes (8, 9), understanding the mechanisms which govern cell shape changes in complex geometrical environments is of particular importance.

Focal adhesions (FAs) represent a key player both for the definition of cell shape and cell mechano-sensing. FAs indeed form a mechanical connection between the acto-myosin cytoskeleton and the ECM, so that the clustering of FA components is necessary for the extension of cell-substrate contact area during spreading (10). On the other hand, FAs respond to the mechanical properties of the substrate through their force-dependent maturation involving the engagement of the integrin ligand, the recruitment of adaptor proteins (such as paxillin or vinculin) (11, 12) and the activation of associated signalling pathways (such as FAK phosphorylation) (8, 13). This process is realized locally in coordination with actin polymerization and myosin II accumulation at the cell leading edge (11, 14). However, the distribution of stresses that FAs collectively transmit to the substrate, as well as the force balance requested at the cell scale, suggest a global regulation of the organization of these structures (15, 16).

To understand this, we need to consider the biological players that govern cell shape changes at the cellular level. One of these players is the acto-myosin cortex, a thin cross-linked network which lies underneath the cell membrane of contact-free cell surfaces (17). A balance between cytosol pressure and an effective surface tension encompassing actin cortex tension and cell adhesion, is then hypothesized to govern global cell shape through its curvature (18–20).

Only a few studies focused on the cooperation between the local clustering of adhesion proteins and three-dimensional cell-scale shape changes. In fully spread cells, it has been demonstrated that circumferential contractile actin arcs, originating from the cell front and linked to FAs, squeeze the cell body and acto-myosin cortex, pointing towards a local control of third-dimensional cell shape (21). On the other hand, in cells spreading between two micro-plates, we have shown that the macroscopic cell body contact angle is a good predictor of cell traction force and the subsequent formation of adhesion complexes (22). Along this line, a phenomenological rule has established a correlation between the distance from the cell edge to the center, traction stress and protrusion rate at the cell edge during cell polarization (23, 24).

Despite these findings, the question of how local and global-scale mechanical processes interplay to define cell shape at the cellular level remains largely unresolved. In this article, we study how three-dimensional cell shape, cell-substrate contact and collective organization of FAs are affected when large scale geometrical parameters of the cell or its environment are modified. We show that the cumulated areas of peripheral FA and cell-substrate contact are conserved whether cells spread confined between parallel plates or unconfined. By changing cell volume, we also demonstrate that these characteristic areas scale with cell size. These effects correlate with changes in cell-substrate contact angle in those conditions, variations which are determined by volume conservation. Finally, we found that allometric relationships between the extent of cell body spreading and lamella size are consistent with the conservation of FA density with respect to cell size and third-dimension confinement. These results demonstrate that the size of cell-substrate contact and collective organization of FAs are regulated by third-dimensional cell-scale geometrical constraints.

## Results

### Cumulated areas of FAs and cell-substrate contact are conserved in confined and unconfined spreading

We analyzed the early spreading of isolated Ref-52 fibroblasts in two different substrate configurations. In the first configuration, the cell is let free to spread on a bi-dimensional glass coverslip. The coverslip is coated with fibronectin to permit integrin engagement. In the second configuration, before the onset of spreading, a glass plate with the same coating is positioned parallel to the coverslip at a distance *h*, with *h* close to the diameter *L*_0_ of the cell while in suspension (*h/L*_0_ = 0.92 ± 0.02, N=15 cells). Thus, cells spread confined in between two parallel plates (Fig. 1-A). Ref-52 fibroblasts expressed a fusion of YFP with paxillin, a marker of FAs. We could thus visualize via TIRF microscopy the evolution of cell-substrate contact (thanks to cytoplasmic paxillin) and the formation of FA-like clusters (intense paxillin aggregates) in the vicinity of the substrate (lower plate only) during spreading in both configurations (Fig. 1-B).

**Fig. 1.**
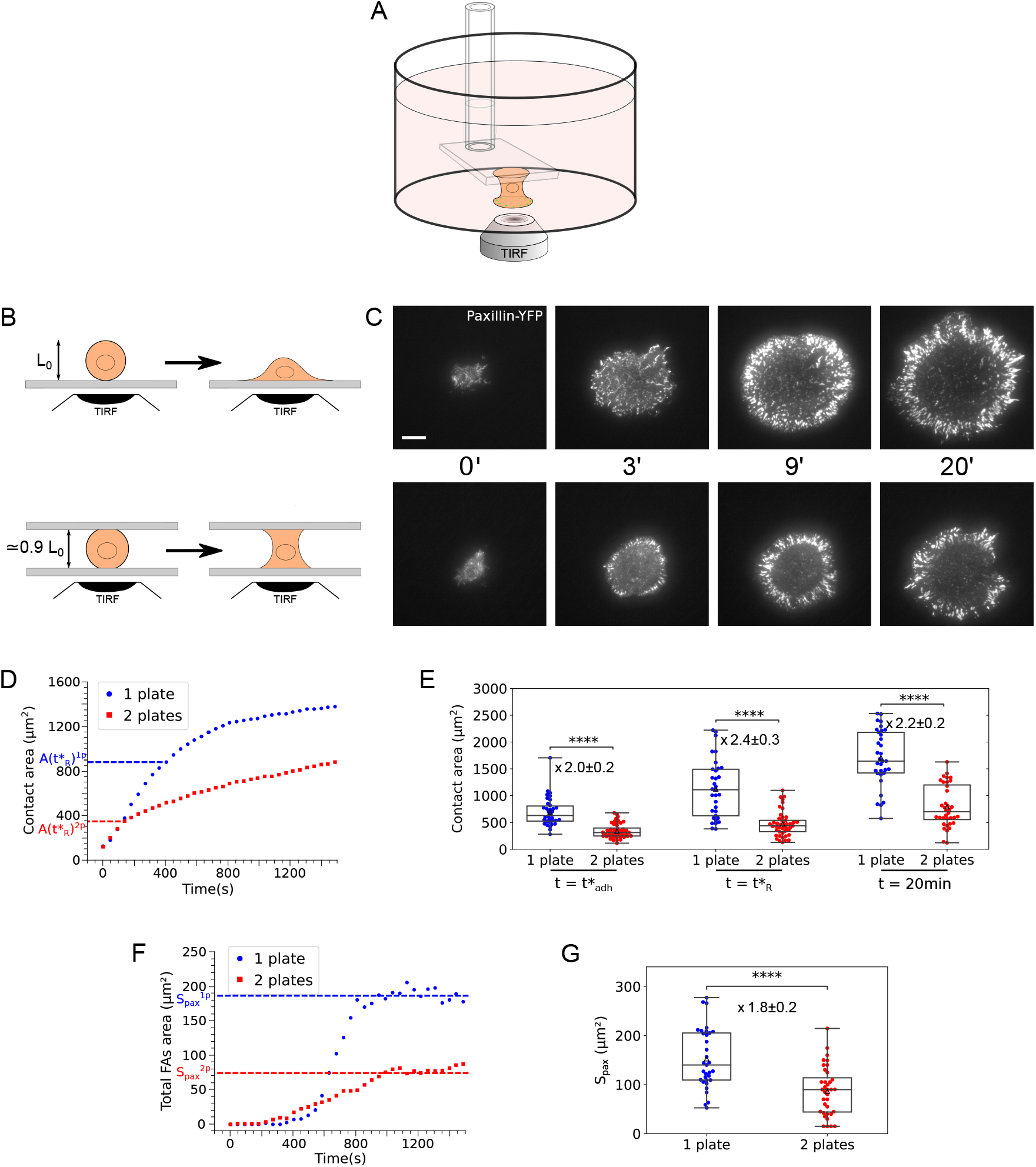
A – Diagram depicting the parallel plate experiment for the analysis of spreading in condition of third-dimension confinement. A microplate parallel to the bottom of the chamber is brought in contact with the cell just after sedimentation and before spreading onset. The contact with the bottom of the chamber is imaged by TIRF microscopy. B – Diagram representing cell spreading configurations for TIRF imaging. Top: A cell in suspension is let free to spread on a uniform bi-dimensional glass substrate. Bottom: A cell is let free to spread between two parallel plates, separated by about 90% of original cell length *L*_0_. C – TIRF imaging of cell-substrate contact of the early spreading of Ref-52 fibroblasts expressing Paxillin-YFP in the two confinement configurations, as described in B. D – Typical evolution of cell-substrate contact area for the single plate (blue) and parallel plates geometry (red). E – Boxplots of characteristic cell-substrate contact areas of cell spreading in the two substrate geometrical configurations. Left: Contact area 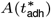 indicating onset of focal adhesions-like paxillin clusters formation. Middle: Contact area 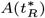 characterizing slow-down of spreading. Right: Contact area *A*(*t* = 20*min*) after 20 minutes of spreading. F – Typical evolution of the integrated area of FA ring for the single plate (blue) and parallel plates geometry (red). G – Boxplot representing the cumulated area of FAs *S*_*pax*_ after 20 minutes of spreading.

In a previous work, we have examined the series of events taking place during spreading in the single plate geometry (22). Briefly, we found that spreading occurs in two phases as previously described (25–27), with a first rapid phase (P1) where cell radius increases proportionally to time (Fig. S1). This phase is followed by a slower phase (P2) after about 450 s of spreading. The slow-down of cell spreading occurs moments after a transition in paxillin organization, with paxillin aggregates evolving from transient dot-like patches uniformly distributed into elongated aggregates, dense in paxillin and analog to FAs, situated at the cell front (Fig. 1-C, top, Supplementary movie 1).

These features are also observable when cells spread confined between parallel plates (Fig. 1-C and 1-D, Supplementary movie 2). First, a ring of paxillin clusters appears after minutes of spreading. Second, the spreading process displays two phases; importantly, the rate of spreading during P1 is unchanged by the addition of the upper plate (*v*^1*p*^ = (3.0 ± 0.2) × 10^−2^µm · s^−1^, N=34; *v*^2*p*^ = (2.7 ± 0.2) × 10^−2^µm · s^−1^, N=45, p-value = 0.33 for two-sided Mann-Whitney test, Fig 1-D and S1-B). However, in the parallel-plate geometry, the transition in spreading regime and peripheral FA ring formation occur at contact areas approximately twice smaller than in the single-plate geometry (factor 2.0 ± 0.2 for the mean contact area at FAs transition 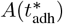; factor 2.4 ± 0.3 for the mean contact area at spreading transition 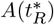, Fig 1-C, 1-D and 1-E). In addition, when spreading has saturated for most cells in both conditions (i.e after 20 minutes), the cell-substrate contact area differs when in confinement, being reduced by a factor 2.2 ± 0.2 in the parallel-plate geometry (Fig 1-C, 1-D and 1-E, and see Fig S1-C for a population representation). It is worth noting that, building on the fact that cells spread similarly on both plates when in a parallel plate geometry (28), the total contact area with the plates is expected to be twice the area measured on the bottom plate here. This implies that the characteristic contact areas of spreading are conserved, irrespective of whether the spreading is in a confined or unconfined geometry.

We next tested whether this accommodation of cell contact area to third-dimensional confinement was accompanied by changes in the surface occupied by FA-like clusters. We found that after saturation of spreading (20 minutes), the contact area occupied by FAs (which we denote *S*_pax_) also adapts to confinement: the total surface occupied by adhesion measured in confined geometry is reduced by a factor 1.8 ± 0.2 as compared to unconfined spreading (Fig 1-C, 1-F and 1-G), suggesting that the cumulated area of FAs on both plates in the confined spreading is equal to the one measured in unconfined (bi-dimensional) spreading. It is important to note that this adaptation of the area of the FA ring occurs while the local environment sensed by the cells at the scale of actin protrusions (glass plates coated with fibronectin) is unchanged between confined and unconfined configurations.

Finally, since cell-generated traction forces are known to adapt to the stiffness of the substrate (28, 29), we investigated, in the parallel plate confined geometry, whether the stiffness of the flexible plate has an impact on the characteristic cell-substrate contact areas (Fig. S2-A). It turns out that, although the maximum cell traction force and the maximum rate of force increase both increase significantly with stiffness (Fig. S2-B and S2-C), stiffness has little or no effect on the evolution of cell-substrate contact and distribution of FAs (Fig. S2-D to S2-H). Therefore, the measurements we present here include data obtained with three different stiffness values (1.5 nN/µm, 12.5 nN/µm and infinite stiffness achieved through feedback-loop control (30)) but we restrict our analysis to the effect of substrate geometry.

### Cell-substrate contact and total FA areas scale with cell size

The above results suggest that cell-scale geometrical constraints determine the overall size of cell-substrate contact and collective organization of FAs. To further test this hypothesis, we varied cell volume and analyzed the evolution of cell-substrate contact on bi-dimensional substrates (Fig. 2-A). Varying cell volume also has physiological relevance since volume varies along cell cycle (31, 32) and according to nutrient conditions (33). We induced the formation of larger cells by culturing them with a low dose of Mitomycin C (MMC), which blocks mitosis without inhibiting protein synthesis (34). On average, before spreading, MMC-treated cells have a 2-fold increase in their apparent surface 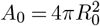, with *R*_0_ the cell radius in suspension, and a 2.8-fold increase in volume (Fig. S3-A to S3-C).

**Fig. 2.**
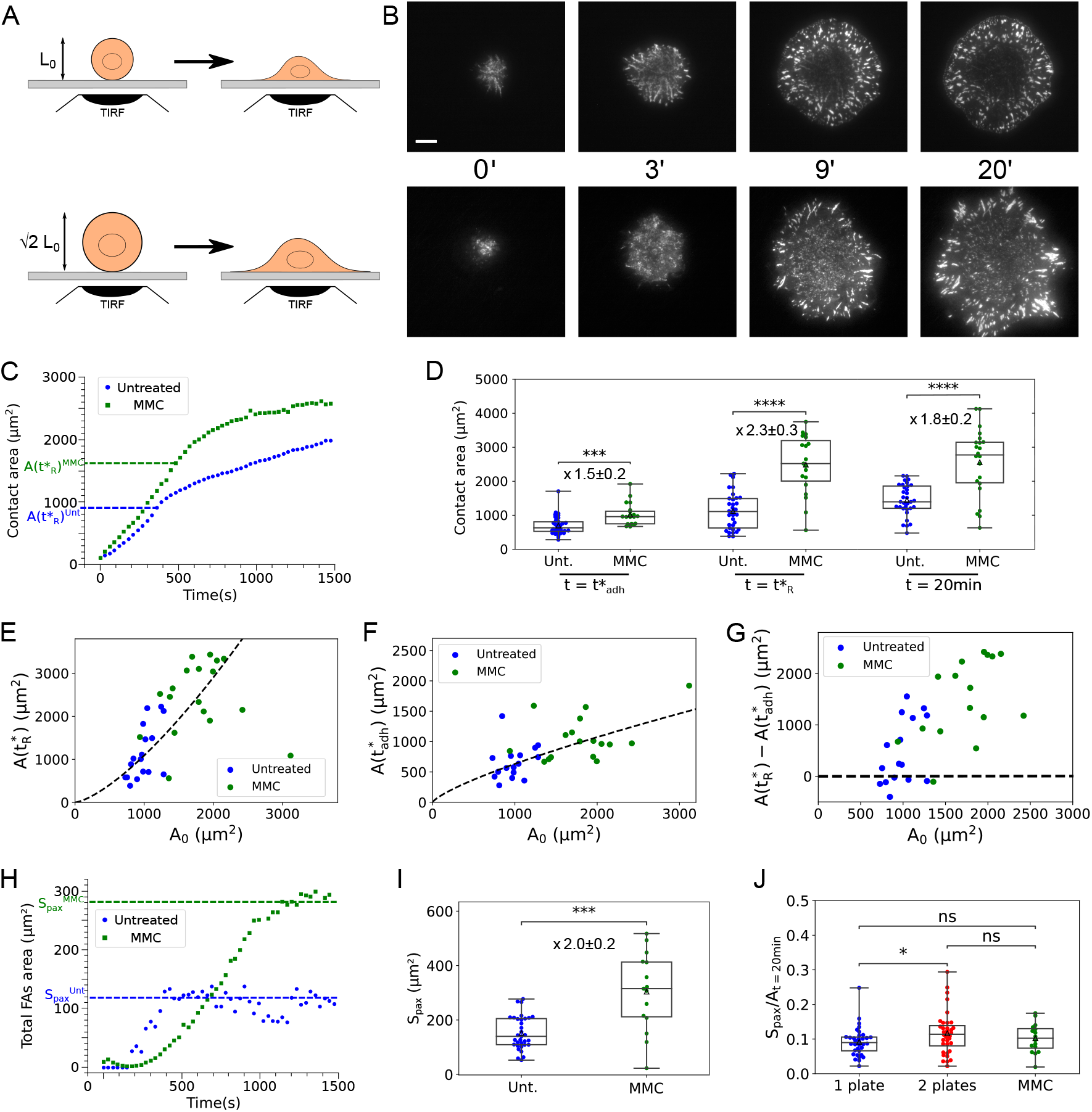
A – Spreading of cells with different size. Top: Untreated cells, normal cell size. Bottom: Cells treated with mitomycin C, large cell size. (Cell volume is increased 2.8 times on average.) B – TIRF imaging of cell-substrate contact of the early spreading of Ref-52 fibroblasts expressing paxillin-YFP in untreated cells (top) and cells treated with mitomycin C (bottom). C – Typical evolution of cell-substrate contact area in untreated (blue) and mitomycin C-treated (green) cells. D – Boxplots of characteristic cell-substrate contact areas of cell spreading in untreated and mitomycin C-treated cells. Left: Contact area 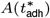 indicating onset of FA-like paxillin clusters formation. Middle: Contact area 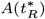 characterizing slow-down of spreading. Right: Contact area *A*(*t* = 20*min*) after 20 minutes of spreading. E – Contact area 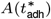 as a function of cell surface in suspension *A*_0_ in untreated (blue) and Mitomycin-C treated cells (green). Dotted line: power-law fit. F – Contact area 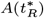 as a function of cell surface in suspension *A*_0_ in untreated (blue) and Mitomycin C treated cells (green). Dotted line: power-law fit. G – Difference between spreading slow-down and onset of focal adhesions-like clusters formation contact areas as a function of cell surface in suspension *A*_0_. H – Typical evolution of the cumulated area of FAs for untreated (blue) and mitomycin C-treated cells (green). I – Boxplot representing the cumulated area of FAs after 20 minutes of spreading for untreated (blue) and mitomycin C-treated cells (green). J – Boxplot representing the ratio of the cumulated area of FAs over cell-substrate contact area for untreated cells spreading on a single plate (1 plate), MMC-treated cells spreading on a single plate (1 plate - MMC) and for untreated cells spreading between two parallel plates (2 plates) after 20 minutes of spreading.

Spreading of MMC-treated cells exhibit the same qualitative characteristics as the one of non-treated cells (Fig. 2-B, 2-C, Fig. S3-D and Supplementary movie 3). Like in the confined geometry, the spreading rate during P1 is not significantly modified by changes in cell size 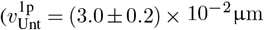 *·* s^−1^, N=34; 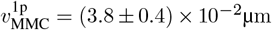 *·* s^−1^, N=21, p-value = 0.11 for two-sided Mann-Whitney test, Fig. 2-C, Fig. S3-E). This is in agreement with the hypothesis that the extension of the lamellipodium is controlled locally by actin polymerization during this phase (22), and that this phase is characterized by non-specific interactions with the substrate (26). However, the areas characterizing spreading are increased in MMC-treated cells compared to untreated cells, by a factor 1.5 ± 0.2 for 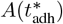, which characterizes the onset of FA maturation, and by a factor 2.3 ± 0.3 for 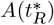, which characterizes the spreading slow-down (Fig. 2-C and 2-D). The dependence of these characteristic areas according to cell size was confirmed when analyzing cells individually with respect to their apparent surface in suspension *A*_0_. We found that both 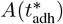 and 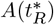 scale with *A*_0_ but with a different exponent: 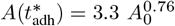 and 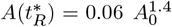 (Fig. 2-E and 2-F). Consequently, the difference in contact area between these two events also increase with cell size (Fig. 2-G). Note also that the exponent 1.4, close to 1.5, suggests that cell contact area at spreading transition scales like cell volume (Fig. S3-F).

Finally, at saturation of spreading, the cumulated area of the peripheral paxillin clusters *S*_pax_ increases following MMC treatment, in proportion with the increase of cell contact area *A*: factor 2.0 ± 0.2 for *S*_pax_(*t*=20min) and 1.8 ± 0.2 for *A*(*t*=20min) (Fig. 2-D, 2-H and 2-I).

### FA areal density is conserved irrespective of cell size and third-dimension confinement

The observation that, at the final stage of spreading, both cell substrate contact area and area of FA ring are rescaled in the same way in confined and unconfined geometry, and whether cells are treated with MMC or not, implies that the ratio *S*_pax_*/A*(*t* =20min) is a conserved quantity. In fact, in all these conditions, the area occupied by the peripheral ring of FAs represents about one tenth of the cell-substrate contact area: 0.09 ± 0.01 for single plate geometry, 0.12 ± 0.01 for parallel plates geometry and 0.10 ± 0.01 for MMC-treated cells (Fig. 2J).

Intra-population comparison over these different conditions confirm that such an adaptation occurs even at the single cell level (Fig. 3-A). A linear regression on the pooled data gives a prefactor equal to 0.09, very close to the mean values of *S*_pax_*/A*(*t* =20min) obtained for each condition. This conservation of the FA areal density is surprising since FAs are mainly distributed along cell perimeter and, therefore, we would rather expect the ratio of cumulated FA area over cell perimeter *S*_pax_*/*(2*πR*(*t* =20min)) to be conserved. Since this is not the case, a non-trivial process must govern the collective organization of FAs and its adaptation to cell-scale geometrical constraints.

**Fig. 3.**
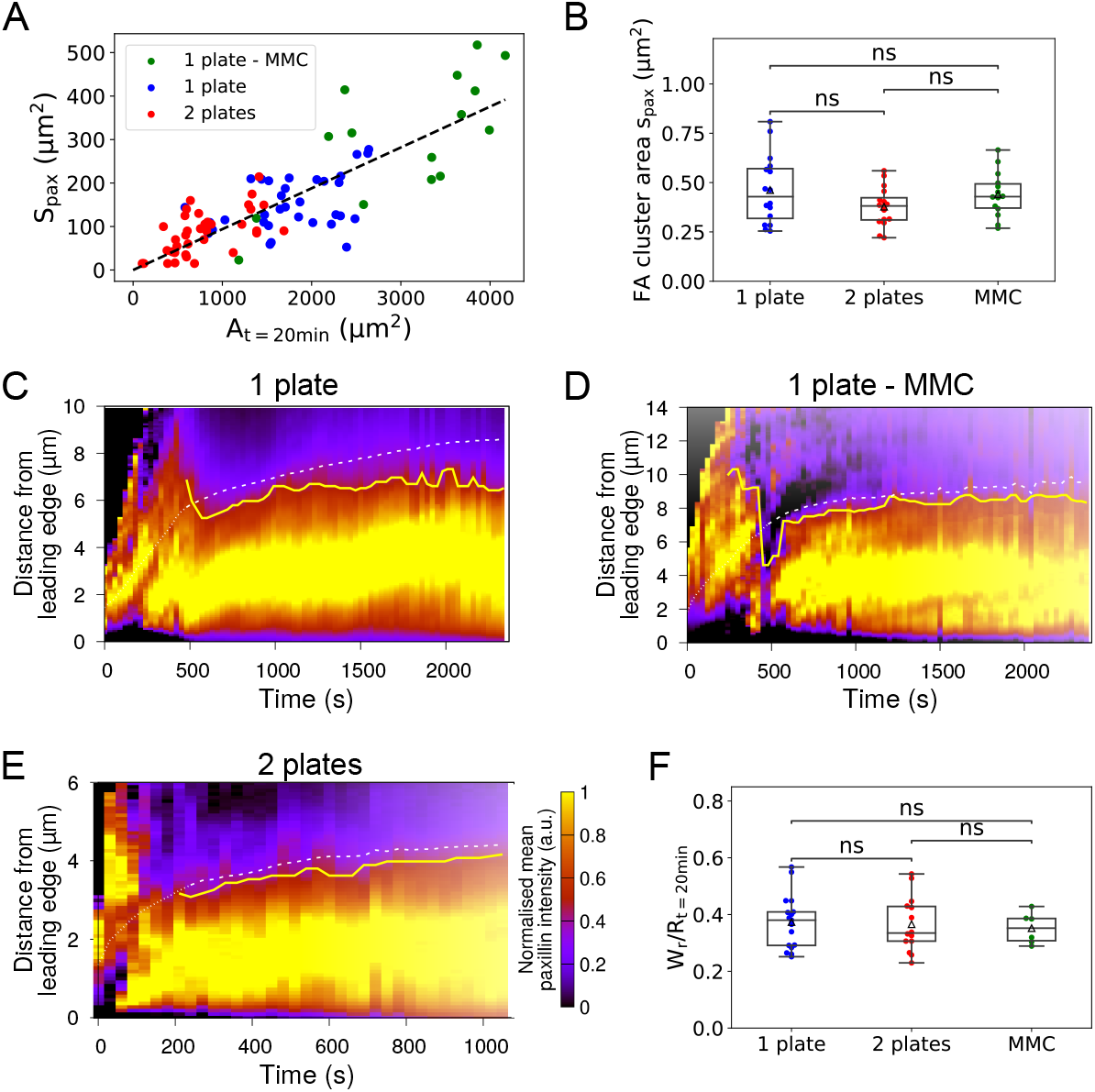
A – Cumulated area of FAs *S*pax as a function of cell contact area *A* after 20 min of spreading for untreated cells spreading on a single plate (1 plate), mitomycin C-treated cells spreading on a single plate (1 plate - MMC) and for untreated cells spreading between two parallel plates (2 plates). B – Individual paxillin clusters area *s*pax. Each point represent the average for an individual cell over N=8 clusters. C - Kymograph of normalized paxillin intensity as a function of the distance to the cell edge and along time for a cell spreading on a single plate. Solid yellow line: paxillin ring width *W*_*r*_. Dashed white line: estimation of *W*_*l*_ from allometry relation *W*_*l*_ = *β/*(1 + *β*)*R* (see Fig. 6). D- Same as Fig. C for a cell treated with mitomycin C and spreading on a single plate. E - Same as Fig. C for a cell spreading between parallel plates. F - Ratio of paxillin ring width over cell contact radius measured after 20 minutes of spreading for untreated cells spreading on a single plate (1 plate), mitomycin C-treated cells spreading on a single plate (1 plate - MMC) and for untreated cells spreading between two parallel plates (2 plates).

We propose a model to comprehend how this cumulated FA area scales like the total contact area, regardless of cell and substrate geometry (Fig. 3-A). In line with other theoretical work (35, 36), we assume that, during early spreading, before the cell polarizes, FAs form at constant density *d*_adh_ along the contact line. Thus, when a new row of FAs forms, the number of FAs (*N*_FA_) varies like:

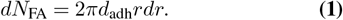

Integrating over the width of the FA ring (which we denote *W*_*r*_), we have:

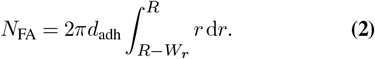

We observed that the area of individual adhesion clusters *s*_pax_ is constant in the different substrate or cell geometries tested (Fig. 3-B). Consequently, the cumulated area of FAs is:

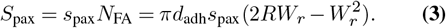

For the ratio *S*_pax_*/A*(*t* = 20*min*) to be constant, we then expect the width of the FA ring *W*_*r*_ to be proportional to the cell contact radius *R*.

To test this, we determined the width of the FA ring *W*_*r*_ by averaging paxillin intensity over cell contour and representing it as a function of the distance to the cell edge at each time-point of the spreading process (see Methods). Figures 3-C, 3-D and 3-E represent the spatio-temporal distribution of paxillin normalized by the maximum intensity along cell radius at each time-point for typical samples in the single plate, parallel plates and MMC-treated conditions respectively. At early time-points, the intensity is homogeneous over the cell-substrate contact, while, during the second phase of spreading, it is higher near the leading edge compared to the cell central region. To determine the position of the inner boundary of the paxilin ring, we used a threshold on the normalized intensity map corresponding to half the maximum intensity at each time-point of the second phase of spreading P2 (yellow lines, Fig. 3-C to 3-E). With this method, we found that the ratio of paxillin ring width over cell contact radius after 20 minutes of spreading *W*_*r*_*/R*(*t* =20min) was independent of substrate geometry or cell size (Fig. 3-F). This scaling was valid not only at 20 minutes but all along the second phase of spreading (Fig. S4-A). According to Equation (3), the scaling between paxillin ring width *W*_*r*_ and cell contact radius R could then explain the conservation of FA areal density, i.e how the surface of FAs *S*_pax_ scales like the contact area *A*(*t* =20min).

In sum, our results point towards a cell-scale mechanism that regulates the size of the paxillin ring along with cell substrate contact radius, while both of these quantities adapt to cell size and third-dimensional confinement.

### The time-evolving 3D body shape of spreading cell scales with cell size and is affected by confinement

Previous work revealed that cell body contact angle, a cellscale parameter, dictates the onset of traction force generation and growth of FAs during spreading between parallel plates (22). Cell-substrate contact angle indeed reflects the equilibrium of tensions between the cell, its substrate and the medium. Therefore, we quantified the effect of cell confinement and cell size on the cell contact angle from profile views of the spreading process imaged in bright-field (Fig. 4-A and 4-B; Supplementary movies 4 & 5).

**Fig. 4.**
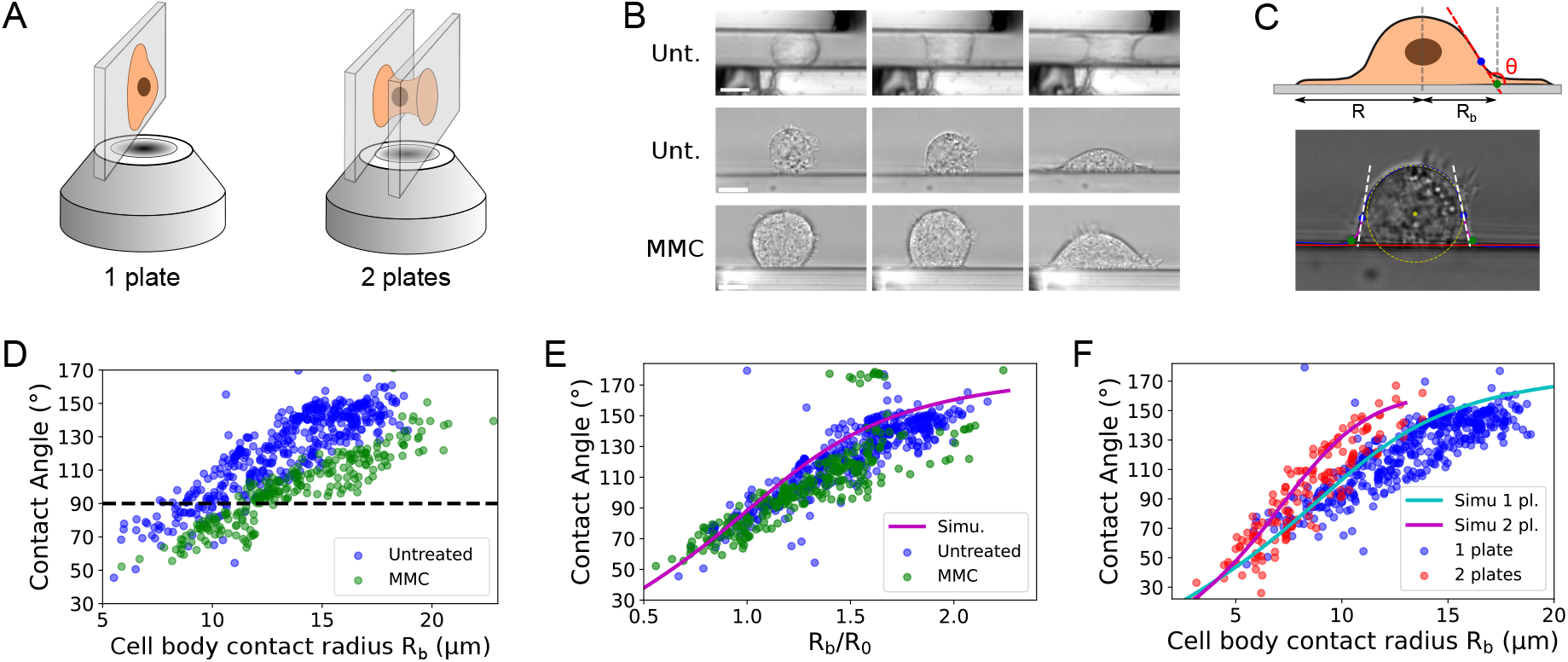
A – Diagram representing imaging configuration of cell spreading imaged in profile. Left: Spreading on a single glass plate, aligned perpendicularly to the objective focal plane. Right: Profile imaging in the parallel plates assay. B – Profile imaging of a cell spreading between parallel plates (top), on a single plate (middle) and a mitomycin C-treated cell spreading on a single plate (bottom). Scale: 10µm. C – Diagram and picture showing the definitions used for cell contact angle and cell body contact radius. D – Cell contact angle *θ* as a function of cortex contact radius *R*_*b*_ for untreated (blue, N=10) and mitomycin C-treated cells (green, N=4). E – Cell contact angle as a function of cell body contact radius *R*_*b*_ normalized by cell radius in suspension *R*_0_. The solid line represents the behavior of a spherical cap of constant volume (see also Fig. 5). F – Cell contact angle *θ* as a function of cell body contact radius *R*_*b*_ for cell spreading on a single plate (blue) or in between parallel plates (red, N=6). The solid lines represents the behaviors of geometrical models as described in the main text (see also Fig. 5).

The automatized segmentation of cell profile and determination of contact angle is described in Methods (Fig. 4-C and Supplementary movie 6). Note that the contact radius measured here — which we name cell body contact radius and denote *R*_*b*_ on the figures because it corresponds to the apparent contact point between the cell body and the plates — does not correspond to the contact radius *R* measured by TIRF microscopy. Indeed, the extremity of the flat lamella at the cell front could not be detected in the profile view experiments realised in bright-field imaging (see Fig. 4-C).

As expected, the cell contact angle (see definition in Fig. 4-C) increased during spreading on a single plate. But, at equivalent contact radii, the cell contact angle was lower in MMC-treated cells compared to non-treated cells throughout spreading (Fig. 4-D). This is the case even for small contact radii corresponding to the first phase of spreading P1, during which the contact radius increases at the same rate in the two conditions (Fig. 2-C). Therefore, there is a discrepancy between the rate at which the contact with the plates and the cell contact angle increase in untreated and MMC-treated cells during the first phase of spreading. Nevertheless, after rescaling the contact radius by the radius of the cell in suspension *R*_0_, the data of contact angle as function of the normalized contact radius collapse on a unique “master” curve (Fig. 4-E). This indicates that cell spreading follows the same dynamics, regardless of cell size or MMC treatment. Notably, those results also indicate that cell volume and cell contact angle cannot be regulated independently.

We next analysed how cell contact angle evolves in the context of cell confinement. Here, the contact angle as a function of contact radius increases with a larger slope when cells are confined between parallel plates than when they are free to spread on a single plate (Fig. 4-F). Like for MMC-treated cells, this effect is observable also during the early phase P1, although the cell-substrate contact areas increased at the same rate in both geometries during this phase of spreading (Fig. 1-D and Fig. S1-B). Thus, contact angle evolution is not only driven by cell-substrate contact interactions, but is also dictated by third-dimension confinement and cell size. Also, it appears that spreading transitions occur at lower contact areas when the contact angle evolves more rapidly. This is consistent with our previous findings showing that the growth of peripheral FAs closely follows the out-of-plane cell traction force, past a specific contact angle (close to 90°) (22).

### Substrate geometry and volume conservation determine three-dimensional cell shape changes during spreading

To understand how cell size and substrate geometry could be at the origin of these differences in threedimensional cell shape changes, we designed a geometrical model of cell spreading (see details in Appendix 1 of Supplementary Materials and Fig. S5-A and S5-B). The modeling idea stems from the fact that cell body shape is known to result from the cortical tension (37), leading to a mechanical balance governed by Laplace law. Making a further assumption of isotropic cortical tension (20, 37, 38) yields that the shape should be of constant mean curvature (CMC). For a given cell volume and for any cell body-substrate contact radius *R*_*b*_(*t*), there is a unique CMC shape, that is, a spherical cap in the unconfined geometry and a portion of either a nodoid or an unduloid in the confined geometry (Fig. S5-C). Assuming that the volume of the cell is conserved throughout spreading, equal to the values shown in Fig. S3-C for untreated and MMC cells respectively, we could thus make predictions of its shape evolution as a function of the cell body contact radius, either in unconfined or confined geometry (Fig. 5, Supplementary movies 7, 8 and 9). In particular, in this model, the data of volume and current cell contact radius are predictive of the contact angle independently of the cortical tension. This is because cell body pressure is expected to adjust in proportion to cortical tension.

**Fig. 5.**
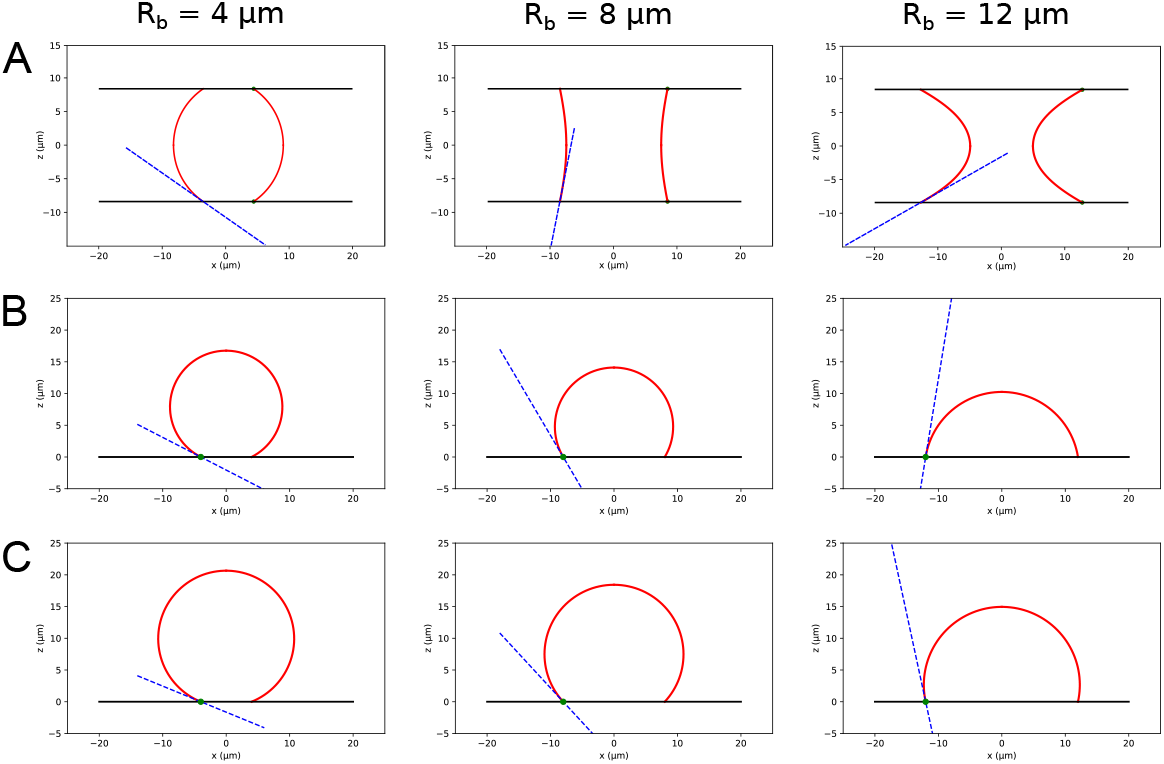
Geometrical modelling of three-dimensional cell shape changes during spreading using constant mean curvature surfaces. A – Cell shape changes of a cell of constant volume spreading between parallel plates (black lines) and whose free edges (red lines) are modelled by a nodoid or unduloid in contact with the plates. Blue dotted lines indicate the tangent to the free edge at the contact point with the plate, defining the contact angle. B – Same as A for a spherical cap (red line) model of constant volume mimicking cell spreading on a bi-dimensional substrate (black line). C – Same as B for a spherical cap of larger initial volume, mimicking spreading under mitomycin C treatment.

In the unconfined geometry, the dorsal arc radius evolves with cell contact radius consistently in the model and in measurements from live cells, indicating that the model can describe global cell shape changes in this configuration (Fig. S5-D). The size-independent evolution of the cell contact angle observed in untreated and MMC-treated cells were also well recapitulated by the geometrical model (Fig. 4-E, solid line). In addition, in the model of parallel plate spreading, the contact angle evolves faster as a function of cell contact radius than in the single plate geometry. Furthermore, the geometrical model reproduces the different rates of contact angle evolution observed in cells spreading on a single plate or in between parallel plates (Fig. 4-F, solid lines). Altogether, this shows that the different rates of three-dimensional cell shape changes, represented by the evolution of cell contact angle, can be explained by two geometrical ingredients acting at the local and global scales respectively: cell-substrate contact expansion, on the one hand, and volume conservation, on the other hand.

The cause leading to a size-dependent relation between cell body contact radius and contact angle is apparent when one considers that the shapes are self-similar when scaling the cell with its size, and that this self-similarity conserves angles whereas distances need to be rescaled. It is less obvious in general to determine how the presence of two plates rather than one only affects the contact angle. However, considering the stage of spreading for which the cell body makes a 90° angle with the substrate, the corresponding shapes of constant mean curvature are respectively a hemisphere and a cylinder of the same volume. In this case, a cylinder will have smaller a basal area, which accounts for the fact that contact angle increases faster in the parallel-plate configuration. Consequently, because of volume conservation, increasing cell confinement is akin to reducing cell size as far as the evolution of the three-dimensional cell shape is concerned. This could explain how those different geometrical perturbations affect the characteristic spreading areas in a similar way.

### The lamella size scales with the cell body contact radius during spreading and coincides with FA ring

On the one hand, we show that the three-dimensional cell shape (quantified via the cell contact angle) adapts to cell size and third-dimensional confinement because of volume constraints; on the other hand, we observed that the width of the FA ring adapts to cell contact radius, regardless of geometrical confinement, and that this permits the conservation of FA areal density. We next wanted to identify a potential link between those cell-scale geometrical relationships in and out of the plane of spreading. Since FAs are found within a flat compartment, called the lamella, present at the leading edge of spreading cells (39) and because the distinction between cell body and lamella is based on their geometrical shape, we went on to quantify the sizes of these respective compartments along spreading. For that, we fixed cells at different time-points during spreading. An F-actin-rich flat compartment was indeed present at the cell leading edge at all stages of spreading (Fig. 6-A). The cell contact radius *R* was thus found to be composed of the cell body contact zone, of radius *R*_*b*_, and the lamella of width *W*_*l*_.

**Fig. 6.**
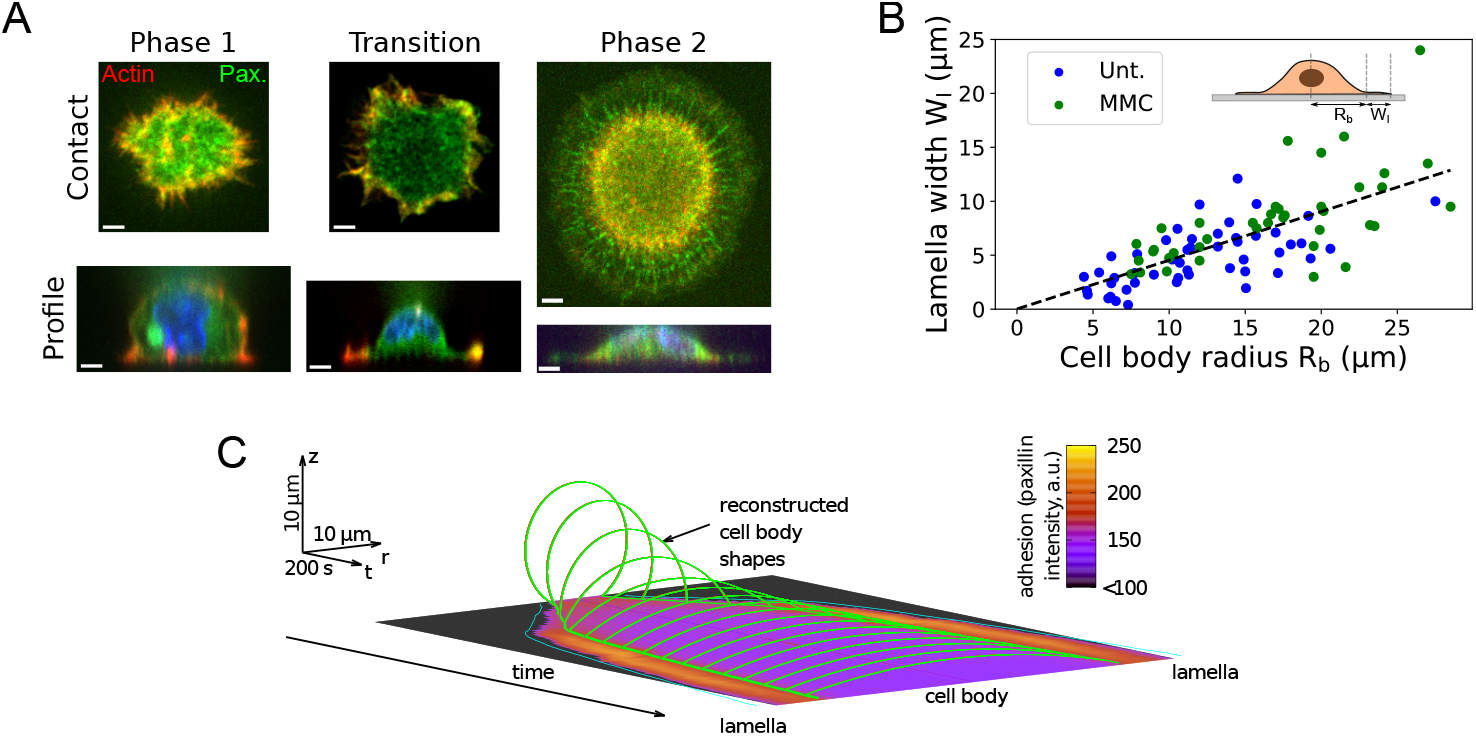
A – Cell shape (profile and cell-substrate contact) imaged through confocal imaging of spreading cells fixed at different stages of spreading. B – Lamella width *W*_*l*_ as a function of cell body contact radius *R*_*b*_ for untreated (blue) and mitomycin C-treated cells (green). C Reconstructed cell body shapes and lamella growth along spreading (for the sample analyzed in Fig. 3C). The averaged spatial distribution of paxillin intensity along time is shown at the cell-substrate interface.

We first verified that cell fixation did not influence threedimensional cell shape by measuring cell contact angle with respect to cell body radius *R*_*b*_ in fixed cells and live cells (Fig. S6-A and S6-B). Second, we found that the size *W*_*l*_ of the lamella increased along spreading, in proportion with the contact radius *R*_*b*_ and regardless of the phase of spreading (Fig. 6-B). Notably, the respective sizes of these two cell compartments (cell body and lamella) followed the same trend in MMC-treated cells, showing that this behavior was independent of cell size and contact angle progression.

Pooling the datapoints from untreated and MMC-treated cells together, a linear regression 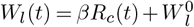 indicated that *β* = 0.45 and 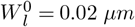 (coefficient of determination *R*^2^ = 0.49). This suggests that the sizes of the lamella and the contact region of the cell body are intrinsically linked. We could then compare the lamella size *W*_*l*_ to the width of the FA ring *W*_*r*_. Note that, whereas the lamella is present and thus *W*_*l*_ is defined at all stages of spreading, the width *W*_*r*_ is defined only when FAs have matured, thus during the second phase of spreading. Considering that *R* = *R*_*b*_ + *W*_*l*_, we find that 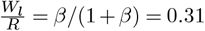, very close to the ratio 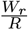 measured during the second phase of spreading (Fig. 3-F), which was equal to 0.37 ± 0.02 for the single plate geometry, 0.37 ± 0.03 for the parallel plate geometry and 0.35 ± 0.02 for the MMC-treated spreading on a single plate. We conclude that, since *W*_*l*_(*t*) ≃ *W*_*r*_(*t*) irrespective of cell size, FAs are located within the lamella during the slow spreading phase P2.

In summary, our data suggest that the growth of FAs is controlled by the cell contact angle, whose evolution depends on cell and substrate three-dimensional geometry. Because the cell-substrate contact area grows at the same rate during P1 independently of these constraints, then FAs start forming at different contact areas. Subsequently, while spreading proceeds, FAs mature and fill the lamella, whose size is proportional to the contact radius (see overlay of cell body reconstruction and lamella size on Fig. 6-C).

## Discussion

At all scales in biology, allometric relationships link the size of the parts or compartments of a system to the size of the entire system. It is the case, for example, for body segments and the size of organisms in tetrapods (40), like for the volume of mitochondria that scale like cell volume in budding yeast (41).

Because fibroblasts evolve in complex and dynamic threedimensional environments within tissues while being also subjected to changes of their own size along the cell cycle, they need to coordinate shape changes occurring at the cell scale (including cell body shape) together with changes in the size of the cell-substrate contact. The mechanical balance implies that cells adapt their shape at the cell scale, but cell shape changes are governed by molecular-scale actors (plasma membrane, cytoskeleton, adhesion complexes). It is still unclear whether the cell shape changes result simply from the integration of local-scale events at the cell scale or whether large-scale phenomena could feed back on the local scale to control the dynamics of molecular actors. This question is particularly debated in the context of cell shape adaptation to changes in the rigidity of their substrate, where mechanisms of adaptation at the cell-scale (governed by the collective dynamics of molecular motors) (29, 42) and at the local scale (through a molecular clutch present in FAs (43)) have been unravelled.

It was previously demonstrated that the formation of FAs is controlled by local traction forces generated by myosin II at the junction between the lamellipodium and the lamella (11, 14). Here we find that the rate of spreading is largely unaffected by cell-scale conditions, both in fast and slow spreading phases, P1 and P2, whereas on the other hand the threshold for a change of behaviour (the switch between P1 and P2) depends on cell-scale phenomena. This indicates that spreading and FA assembly are not purely kinetic processes as previously suggested (44). Various hypotheses have been proposed so far that could potentially explain how a cell scale parameter could control the formation of FAs at the local scale.

First, theoretical work suggested that an increase in the length of the contact line could trigger FA clustering via a phase transition process relying on short-range protein bridging (36). In our system, FAs should then form at a conserved size of individual cell-substrate contacts irrespective of the substrate geometry (single plate or two parallel plates). However, our data show that FA assembly is rather triggered when the total cell-substrate area reaches a threshold that is proportional to the cell size.

Another mechanism of transition was proposed that would rely on a burst of plasma membrane tension after exhaustion of a pool of free membrane. That event could trigger the transition between the two phases of spreading, and subsequent growth of FAs (45). This is consistent with a spreading transition occurring at a conserved total cell-environment contact area, as observed here, rather than at a given size of individual contacts. However, although a sudden membrane flattening accompanies the transition from fast to slow cell spreading, these two events occur after the onset of FA formation (22). This sequence of events is confirmed by our data and even more compelling in the case of larger cells. Conversely, in some rare events (n/N=3/30) where spontaneous disassembly of a first ring of FAs happened after entering phase P2, it was correlated with an acceleration of cell spreading at a rate similar to P1, before FAs re-formed into a second, larger peripheral ring (Fig. S7 & Supplementary movie 10). These observations thus rather suggest that FA growth is the cause rather than consequence of the transition.

Finally, a third mechanism that could trigger the formation of FAs is a change in the force balance at the cell scale, whereby the (possibly unchanged) actomyosin cortical tension which is internally balanced at the beginning of cell spreading starts being transferred to the substrate at some stage. In (22), we show that indeed FA formation coincides with the stage where the cell—substrate apparent contact angle reaches 90°, an angle beyond which internal balance of cortical tension becomes geometrically impossible.

Although the trigger of FA growth and maturation is a global-scale mechanism, their individual growth has all the features of a local phenomenon: in all conditions, we find that individual FAs have the same area. Consistently with (35, 36), their density appears to be constant. Finally, there is no apparent variability in the intensity of each FA, suggesting that they carry the same adhesion energy. Taken together, these three local and invariant intensive characteristics (density, individual size and intensity) lead to an invariant adhesive surface – and presumably adhesive energy – per unit area of the zone over which they are distributed. Such a conserved invariant quantity can thus be explained by purely local mechanisms. However, we find that, on the other hand, the total adhesive surface (cumulated FA area) scales with the total area of contact with the substrate, with a scaling parameter which is robust to the geometrical changes induced by confinement and cell size. Notably, the same effect has been observed on human foreskin fibroblasts confined on bi-dimensional adhesive islands of circular shapes. In that case, the peripheral FA ring represented about 15% of the contact area (46), for fibroblasts here we find that it is approximately 10% of the contact area. Maintaining such a scaling irrespective of whether the cell is confined or not requires a global regulation. Thanks to 3D reconstruction of the geometry of the actin cortex, we determine the relative size of the lamella of cells at different stages of spreading. Surprisingly, we show that throughout the different phases of spreading, the lamella length scales with the spread radius of the cell body while they both undergo a more than 5-fold increase, thus being globally regulated whereas the leading edge protrusivity seems to be locally regulated. In order to coordinate the position of the body–lamella transition point, there is thus also the requirement for a global coordination mechanism able to conserve this observed allometry in coordination with leading edge progress. Future work could focus on the origin of this coordination.

## Methods

### Cell culture and mitomycin C treatment

Rat embryonic fibroblasts (Ref-52) stably expressing paxillin-YFP were grown using DMEM supplemented with 10% (vol/vol) heat-inactivated FCS, 2 mM glutamin, 50 units/mL penicillin, and 50 g/mL streptomycin. All cultures were maintained at 37°C under humidified 5% CO2 atmosphere. To increase cell volume, Mitomycin-C (Sigma, M4287) at a concentration of 0.25*µ*M was introduced into culture medium for 2 days prior to experiments. This concentration blocked cell cycle while limiting cell death.

### Spreading assay and TIRF imaging

Before experiments, cells were trypsinized, and maintained under smooth agitation for at least 2 h at 37 °C in medium containing DMEM without Phenol red, supplemented with 10% FBS and 15µL/mL Hepes. Cells were then released in an imaging chamber containing the same medium as during incubation maintained at 37°C. TIRF imaging of spreading was performed every 30s through a PlanApo 100X/1.45 oil objective and a F-View CCD camera (Olympus) using an IX-71 Olympus microscope coupled to a monochromatic (GFP, 450–480 nm) TIRF arm. For the parallel plate assay, a glass microplate was brought above the cell just before onset of spreading thanks to piezo-electric translation stages interfaced on a 3-axis micro-manipulator (PhysikeInstrumente). The glass micro-plate was pulled from rectangular plates using a micropipette puller (PB-7; Narishige) and adjusted with a microforge (MF-900; Narishige); they were then attached to a bent glass capillary and connected to a micro-manipulation apparatus prior to experiment. The bottom of the imaging chamber and the glass micro-plate were incubated in DMEM containing 5 µg/mL fibronectin (F1141; Sigma-Aldrich) prior to experiment to promote specific cell adhesion.

### Spreading assay and profile imaging

For profile imaging of spreading in bright-field, the cells were prepared as described above. For the parallel plates assay, two glass microplates fabricated and coated as described above were positioned at the bottom of the chamber parallel to each other and brought in contact with the cell before to be lifted. For spreading on a single plate, the same micro-plate was aligned perpendicularly to the bottom of the imaging chamber and positioned in contact with the cell. To avoid cell spreading at the bottom of the imaging chamber, a thin layer of PDMS (about 10 µm) was spin-coated on the glass coverslip and coated with Pluronic (1% in water; F127TM; Sigma-Aldrich) for 1h after plasma cleaning.

Spreading was imaged every 10s through bright-field imaging using a CMOS camera (ORCA-Flash 4.0, Hamamatsu) and a 60X water objective (UPLSAPO, Olympus) mounted on an Olympus IX81 inverted microscope.

### Measurement and characterization of cell contact area

Methods for segmentation and transitions in cell-substrate contact area and paxillin organization were thoroughly described in (22). Briefly, for the determination of the cell–substrate contact area from TIRF images of paxillin-YFP, an ImageJ custom-built macro was used. Noise was removed using the functions: *Despeckle, Smooth, Subtract Background* before binarization of the image through a threshold in intensity manually defined. To include some potentially isolated parts, we then used the closure algorithm *Close-*. The contact area 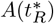 corresponding to the transition in spreading regimes was defined as the largest contact radius 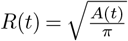 before *R*(*t*) departed from a linear trend.

### Segmentation of FA clusters

The regions covered with FA clusters were defined according to a threshold in intensity from TIRF images of paxillin-YFP. For that, the entropy-based algorithm *RenyiEntropy* (47) based on the entropic analysis of the intensity histogram of the image and implemented in ImageJ was used. During the early phase of spreading 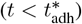, the algorithm detected a region corresponding to the entire contact area, indicating that no region of the contact was specifically intense. We noted 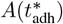, the contact area retrieved from the last frame devoid of adhesion clusters. Then, we noted *S*_pax_ the area of the peripheral ring of intense paxillin-YFP clusters measured after 20 minutes of spreading.

Individual FA clusters were segmented manually. A minimum of 8 clusters per cell were selected based on their aspect ratio (which was chosen superior to 1.8) and on their location (within FA ring). The average FA area was then calculated for each cell tested.

### Determination of the width of the FA ring

To measure mean paxillin intensity with respect to the distance to cell edge, we use the *Erosion* algorithm of *ImageJ*. Because the erosion in ImageJ is performed with a square 3 *×* 3 structuring element, the distance from the leading edge of the *n*-th concentric region is best approximated as 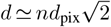 where *d*_pix_ is the pixel size. Next, we plotted the mean fluorescence intensity *I*_raw_(*d, t*) of these concentric regions as a kymograph. In order to detect the distance from which the signal significantly decreases, we first normalized the signal *I*_raw_ by an affine mapping, 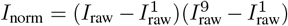 where the 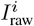 are the *i*-th decile value of *I*_raw_. Next we detected the distance from edge 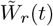 where *I*_norm_ crossed the value 0.5 for the first time while decreasing,

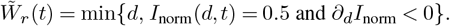

Since occasional large variations are possible over a single frame, we applied a median filter with a window size of 3 in order to define *w*_*r*_,

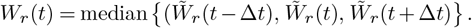

### Segmentation of cell profile and measurement of contact angle

To measure the shape changes of cells profile during spreading, an ad-hoc python algorithm was developed. First, to make the glass plate appear horizontal on the image, a rotation of the image is applied (using a linear Hough transform to detect the glass edge original angle). Optical aberration generated by the glass plate in the cell region is then removed by subtracting to each vertical line of the image the typical profile of the shadow computed from a region of the image free of cell. Finally, the profile of the cell is segmented using the active contour method (48) implemented in the *scikit-image* library (49). After a manual initialization of the cell contour on the first frame, the contour is re-adjusted in each successive image to the new cell shape using the active contour method. These contours are analyzed to determine the geometrical descriptors of the cell shape on each frame (see Fig. 3C). The dorsal arc radius is extracted by fitting an arc of circle to the upper part (representing 2/3 of cell height) of the cell body. The cell body contact diameter is calculated from the two points of the contour on each side of the cell reaching a distance to the plate ≤ 0,75µm. Finally, the macroscopic cell contact angle is determined from the tangent to the contour at the point of inversion of cell contour curvature. For that, the local curvature of the cell contour *κ* was computed along the contour expressed parametrically as 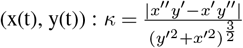. The tangent of the cell contour around this position is subsequently obtained from a linear fit along a 1 *µ*m contour length.

### Confocal imaging

For confocal imaging of fixed cells at various stages of spreading, cells were fixed with paraformaldehyde (15 min; 3% in PBS), permeabilized in Triton X-100 (15 min; 0.2% in PBS), and put in presence of BSA (30 min; 1% in PBS). For actin staining, cells were incubated with fluorescent phalloidin from Fluoprobes (30 min; 0.1% in PBS). Cells were imaged using a confocal spinning disk from Andor Technology coupled to an Olympus IX-81 microscope with a 60X oil objective.

The lamella width and cell body radius were measured based on the same criterion as for the determination of the cell body radius in profile bright-field imaging of spreading: the extremities of the cell body diameter were defined from the two points of the contour on each side of the cell reaching a distance to the substrate ≤ 0,75µm. The lamella width *W*_*l*_ was then calculated from the total cell contact area A measured independently and through the equation *A* = *π*(*W*_*l*_ + *R*_*b*_)^2^, with *R*_*b*_ the cell body radius.

## AUTHOR CONTRIBUTIONS

C. Bimbard performed TIRF measurement and analysis of MMC treated cells; A-A Wahhod and J. Etienne designed and realized spreading simulations; J. Vermeil and R. Bousquet designed the segmentation algorithm for profile view analysis; A. Richert performed confocal imaging; P. Durand and S. Asnacios performed cell profile imaging in confined geometry; D. Pereira and D. Mitrossilis performed image analysis and provided analysis insights; J. Etienne realized FA ring width analysis and wrote the manuscript; A. Asnacios designed the research and wrote the manuscript; J. Fouchard performed all the other measurements and analysis, coordinated the research and wrote the manuscript.

## ACKNOWLEDGEMENTS

AAW was funded by doctoral school IMEP2 of University Grenoble Alpes. DM was founded by the Hellenic Foundation for Research and Innovation (H.F.R.I. 2nd call for the support of Postdoctoral Researchers, project number 01281, FA 94-2/14.10.2020). The study was partially supported by the labex “Who AM I?”, labex ANR-11-LABX-0071, as well as the Université Paris Cité, Idex ANR-18-IDEX-0001, funded by the French Government through its “Investments for the Future” program and also by the Agence Nationale de la Recherche through the project “scEm-bryoMech” ANR-21-CE13-0046. PD, JE, AA and JF are members of GDR 2108 “Approches quantitatives du vivant” of CNRS. JE is member of GDR 3570 MecaBio of CNRS. The authors thank Tom Wyatt, Estelle Hirsinger, Joseph d’Alessandro, Mathieu Hautefeuille and Marc-Antoine Fardin for fruitful discussions and critical reading of the manuscript, and Sophie Gournet for artwork.

