## Supplementary notes and figures for "Allometric spreading and focal adhesion collective organization are coordinated by cell-scale geometrical constraints"

### Appendix 1 : Geometrical description of cell spreading

#### 1. Model

The simplest model of cell body shape corresponds to assume that it is governed only by cortical actomyosin tension and plasma membrane-regulated volume. This has been shown to lead to consistent predictions during cell spreading on two plates (1). Volume regulation implies a pressure difference across the membrane (and the thin actomyosin cortex apposed to it), this pressure equilibrates spatially at a timescale much shorter than spreading (2). Following (3) we further assume that the cortex is under a uniform isotropic tension $\sigma(t)$ and axial symmetry of the shape. The shape of the cell cortex in those conditions can be parameterised as $\Gamma = \{r(z,t), z_{\min} \le z \le z_{\max}\}$ and the volume of the cell body can be calculated by:

$$V(t) = \pi \int_{z_{\min}}^{z_{\max}} r(z,t)^2 \mathrm{d}z$$

It obeys the Young-Laplace equation:

$$H(t) = \frac{r''}{\left(1+r'^2\right)^{3/2}} - \frac{1}{r\sqrt{1+r'^2}} = \frac{\Delta p}{\sigma} \qquad [1]$$

where $r' = \mathrm{d}r/\mathrm{z}$, $\Delta p(t)$ is the pressure difference between the interior and exterior and we define $H$ the mean curvature. Given uniform pressure and tension, $H$ is uniform in space and thus the shape has constant mean curvature and can be a portion of sphere, cylinder, unduloid or nodoid (Fig. S5-C).
For single plate spreading, the only appropriate shape is obviously a spherical cap and it can be readily shown that for a given volume $V$ and basal radius $R_b$ there is only one solution to Eq 1. The ratio $\frac{\Delta p}{\sigma}$ of pressure difference and tension is thus deduced from the volume.
For the spreading between two plates, the shape can be either of a nodoid, unduloid or cylinder depending on the respective values of $R_b$ and $V$. Similar to the single plate case, being given these two values, a unique shape and the corresponding ratio $\frac{\Delta p}{\sigma}$ of pressure difference and tension can be deduced.
In what follows, we reproduce the sequence of shapes of a cell during spreading, either in unconfined (single plate) or confined (parallel plates) geometry, by assuming that it maintains a constant volume $V = V_0$ and varying its radius $R_b(t)$.

#### 2. Single plate geometry: spherical cap

Given $R_b(t)$ and $V_0$, it can be easily verified that spheres of parametric equation

$$\begin{cases} z(s) = R_{cap}(1-\cos(s)) + z_0 \\ r(s) = R_{cap}\sin(s) \end{cases}$$

for a curvilinear coordinate $s \in [0, s_f]$, with $z_0(t) = \pm\sqrt{R_{cap} - R_b}$ and $s_f = \arccos\left(\frac{R_{cap}+z_0}{R_{cap}}\right)$ are solutions. We calculate the radius of curvature $R_{cap}(t) = 2/H(t)$ such that the spherical cap volume $V(t) = \frac{\pi}{6}(R_{cap}+z_0)(3R_b^2 + (R_{cap}+z_0)^2)$ is equal to $V_0$.
The (outer) contact angle at the plate can be calculated as $\cot\theta = -\frac{\mathrm{d}r}{\mathrm{d}z}$, resulting in $\theta = \cos^{-1}(z_0/R_{cap})$.

#### 3. Two plates geometry

When spreading between two parallel plates spaced by height $h$, we assume that the shape is symmetric with respect to the plane $z = 0$ parallel to the plates $z = \pm h/2$ and at equal distance from them. Thus the surface forms a right angle with this plane, $r'(z=0) = 0$. The additional conditions are that $r(z = h/2) = R_b(t)$ and, as above, volume $V(t) = V_0$ allows to set $H(t)$. We proceed with an additional intermediate unknown, $r(z=0,t) = R_e(t)$: for a given $t$, we initialise $R_e(t)$ to a first guess $R_e^0(t)$ and use a fixed point algorithm on $R_e^i$ to reach $V(t) = V_0$.
The problem of integration of nonlinear Eq 1 is thus done with boundary conditions:

$$\begin{cases} r(s_i,t) = R_e(t), & z(s_i) = 0 \\ r(s_f) = R_b(t), & z(s_f) = h/2 \\ \frac{\mathrm{d}r}{\mathrm{d}z}(s_i) = 0 \end{cases} \qquad [2]$$

It is known (4) that depending on $R_e$, the solution can belong to five families of curves: nodoids for $R_e^2 < R_b^2 + (h/2)^2$, a sphere when $R_e^2 = R_b^2 + (h/2)^2$, an unduloid for intermediate values of $R_e$ (but a cylinder for $R_e = R_b$), a catenoid when $R_e = R_e^*$ such that $R_e^* \cosh(h/(2R_e^*)) = R_b$ and nodoids again when $R_e > R_e^*$. In practice, for the range of $R_b(t)$ that is useful to compare with experiments, we find that $R_e < R_e^*$. Note that spheres, cylinders and catenoids can be seen as limiting cases of unduloids or nodoids.

**A. Unduloid.** Its parametric representation (5) is:

$$\begin{cases} z(s) = \frac{b^2}{a}\int_0^s I_u(u)\mathrm{d}u + z_0, \\ r(s) = b\sqrt{\frac{1-\xi_u\cos s}{1+\xi_u\cos s}}. \end{cases} \qquad [3]$$

where

$$I_u(u) = \frac{1}{(1+\xi_u\cos u)\sqrt{1-\xi_u^2\cos^2 u}},$$

and $s$ is a curvilinear coordinate varying in a range $[s_i, s_f]$ which is to be determined, whereas $a$, $b$ and $\xi_u = \sqrt{1 - \frac{b^2}{a^2}}$ are positive parameters also to be determined. It can be shown that such curves obey the equation:

$$1 + \left(\frac{dr}{dz}\right)^2 = \frac{4a^2 r^2}{\left(r^2 + b^2\right)^2} \quad [4]$$

which in turn verify Eq 1 with $H = \pm 1/(2a)$.
Because $r(s_i) = R_e$ is the minimum of $r$ if $R_e < R_b$ (resp. the maximum else), $s_i = 0$ (resp. $s_i = \pi$). Rewriting $b = a\sqrt{1 - \xi_u^2}$ and using the boundary condition $r(s_f) = R_b$, the parameters $b$, $s_i$ and $s_f$ can be expressed as functions of $\xi_u$ and $a$. The additional boundary condition $r'(z = 0) = 0$ injected in Eq 4, yields two possible values of $a$ as a function of $\xi_u$,

$$a(\xi_u) = \frac{R_e(1 - \chi \xi_u)}{1 - \xi_u^2}$$

where $\chi = \pm 1$. It can be shown that $\chi$ has to have the same sign as $R_e - b$ (6), which has the same as the sign as $R_e - R_b$. The remaining parameter, $\xi_u$, can be determined from the implicit equation

$$h/2 = z(s_f(\xi_u, R_b)) = \frac{b(\xi_u)^2}{a(\xi_u)} \int_{s_i}^{s_f(\xi_u, R_b)} I_u(\xi_u, s)\mathrm{d}s \quad [5]$$

As above, the angle at the bottom plate can be found using $\cot\theta = -\frac{\mathrm{d}r}{\mathrm{d}z}(-s_f)$.

**B. Nodoid.** Its parametric representation (5) is:

$$\begin{cases} z(s) = \dfrac{b^2}{a} \displaystyle\int_0^s I_n(u)\,\mathrm{d}u + z_0, \\ r(s) = b\sqrt{\dfrac{\xi_n - \cos s}{\xi_n + \cos s}}. \end{cases} \quad [6]$$

where

$$I_n(u) = \frac{\cos u}{(\xi_n + \cos u)\sqrt{\xi_n^2 - \cos^2 u}},$$

and $s \in [s_i, s_f]$, a range to be determined. The positive parameters $a$, $b$, and $\xi_n = \sqrt{1 + \frac{b^2}{a^2}}$ obey the governing differential equation:

$$1 + \left(\frac{\mathrm{d}r}{\mathrm{d}z}\right)^2 = \frac{4a^2 r^2}{\left(r^2 - b^2\right)^2}. \quad [7]$$

Similarly to the unduloid case, and using the boundary condition at $z = 0$, $a$ can again be expressed as a function of $\xi_n$:

$$a(\xi_n) = \frac{R_e(\chi - \xi_n)}{1 - \xi_n^2}.$$

The remaining parameter, $\xi_n$, is determined by solving the implicit equation:

$$h/2 = z(s_f(\xi_n, R_b)) = \frac{b(\xi_n)^2}{a(\xi_n)} \int_{s_i}^{s_f(\xi_n, R_b)} I_n(\xi_n, s)\,\mathrm{d}s, \quad [8]$$

where the integration bounds $s_i$ and $s_f$ are defined based on the boundary conditions.

**C. Implementation and upper limit.** In Figure 5, we set $V_0 = \frac{4}{3}\pi R_0^3$, where $R_0 = 8.83\mu$m for untreated cells, and $h = 1.9R_0$, in accordance with the experimental conditions. For $R_b$ varying over the range of experimentally observed values, we then calculate the shape of spherical caps of volume $V_0$, and of either unduloids or nodoids of volume $V_0$ and height $h$, as detailed above. We can then deduce the angle that these shape make with the substrate.
There is a lower bound for the volume enclosed in a CMC surface of rotational symmetry having radius $r = R_b$ at $z = h/2$, however to the best of our knowledge there is no general formula allowing to compute it. In practice, we find that solutions cease to exist for our choice of parameters $V_0$ and $h$ beyond a radius $R_b^* \simeq 13\mu$m. There does not exist, e.g., a section of catenoid of height $h$ that has volume $V_0$. The physical understanding of this geometrical limitation is that the model must cease to be valid for cells observed with $R_b > R_b^*$: it is e.g. possible that the symmetries assumed here (axial symmetry, isotropy of the tension) cease to be a fair approximation.

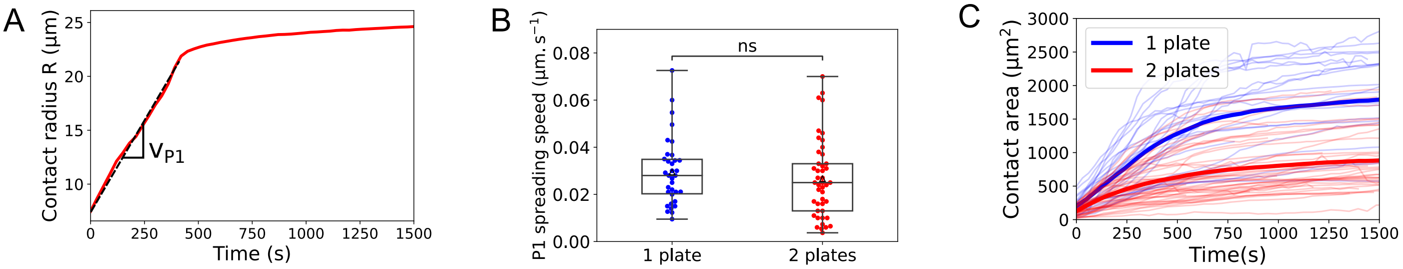


**Fig.S 1.** A - Cell-substrate contact radius as a function of time of a typical Ref-52 fibroblast spreading on a bi-dimensional substrate. $v_{P1}$ describes the spreading speed during the first phase of spreading. B - Boxplot representing the spreading speed during the first phase of spreading P1 in the single plate and parallel plates geometry. C - Contact area as a function of time for all samples tested in the single plate (blue) and parallel plates (red) geometry. Bold lines represent the mean value calculated over all samples at each time-point.

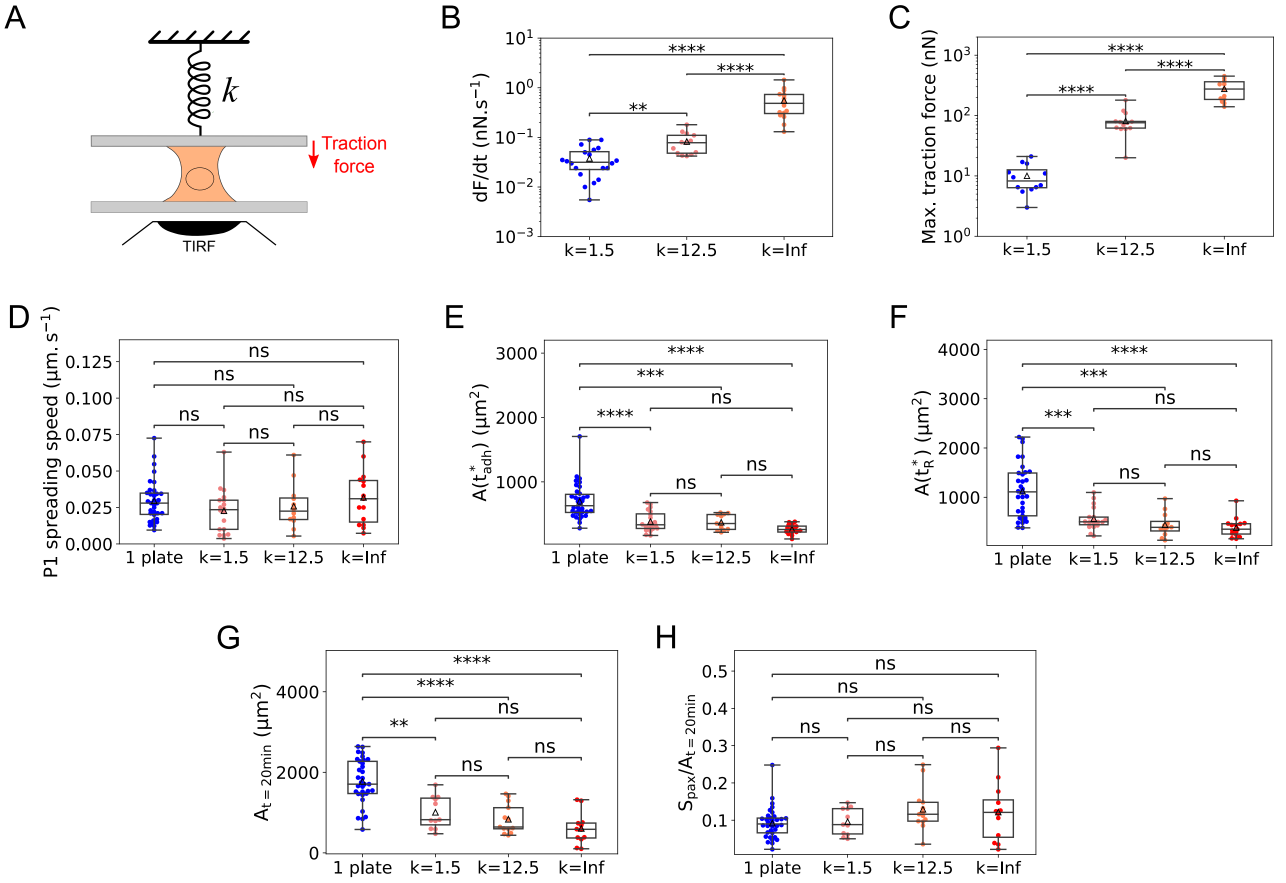


**Fig.S 2.** Effect of upper plate stiffness on cell traction, spreading and focal adhesions formation (k=1.5 nN/µm, N=17; k=12.5 nN/µm, N=11 and infinite stiffness, N=14) and comparison with single plate experiment ('1 plate'). A - Sketch of parallel plate setup. The upper plate is flexible, with stiffness $k$. TIRF imaging is performed at the bottom, rigid plate. B - Maximum rate of traction force increase. C - Maximum traction force. D - Spreading rate during the first phase of spreading (P1). E - Contact area $A(t^*_{adh})$ at which paxillin starts forming aggregates. F - Contact area $A(t^*_R)$ at transition between spreading phases. G- Cell-substrate contact area after 20 minutes of spreading. H - Ratio of focal adhesions ring area over cell-substrate contact area after 20 minutes of spreading.

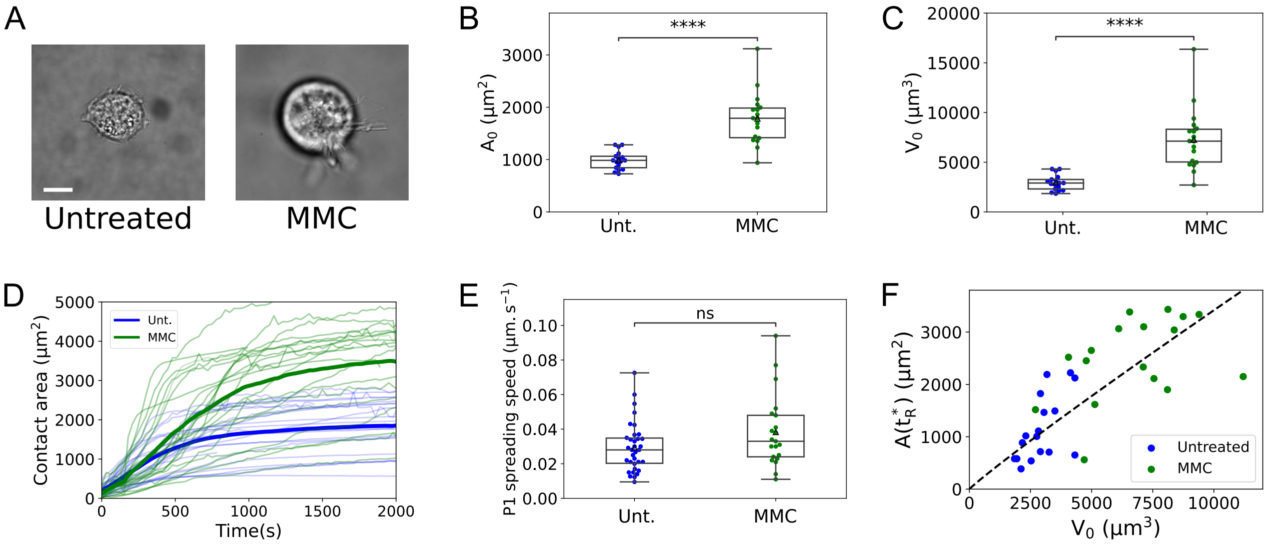


**Fig.S 3.** Effect of mitomycin C treatment (0.25µM, 2 days) on cell size in suspension. A - Bright-field images of Ref-52 fibroblast in suspension treated (MMC, right) or not (Untreated, left) with mitomycin C. Scale bar: 10µm. B - Effect of mitomycin C treatment on cell surface in suspension. C - Effect of mitomycin C treatment on cell volume in suspension. D - Contact area as a function of time for all untreated (Unt., blue) and mitomycin C treated (MMC, green) samples. Bold lines represent the mean value calculated over all samples at each time-point. E - Boxplot representing the spreading speed during the first phase of spreading P1 for untreated and mitomycin C-treated samples. F - Contact area $A(t_R^*)$ as a function of cell volume in suspension $V_0$ in untreated (blue) and Mitomycin C treated cells (green). Dotted line: power-law fit of exponent 0.94, close to 1.

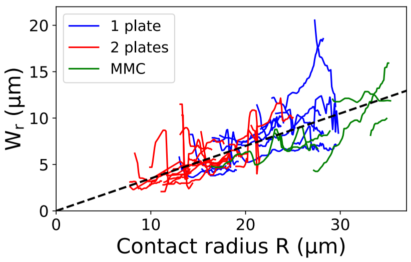


**Fig.S 4.** Width of the focal adhesions ring versus contact radius along the second phase of spreading P2 for the single plate (blue), parallel plates (red) and MMC treated cells spreading on a single plate (green). Each line represent one sample.

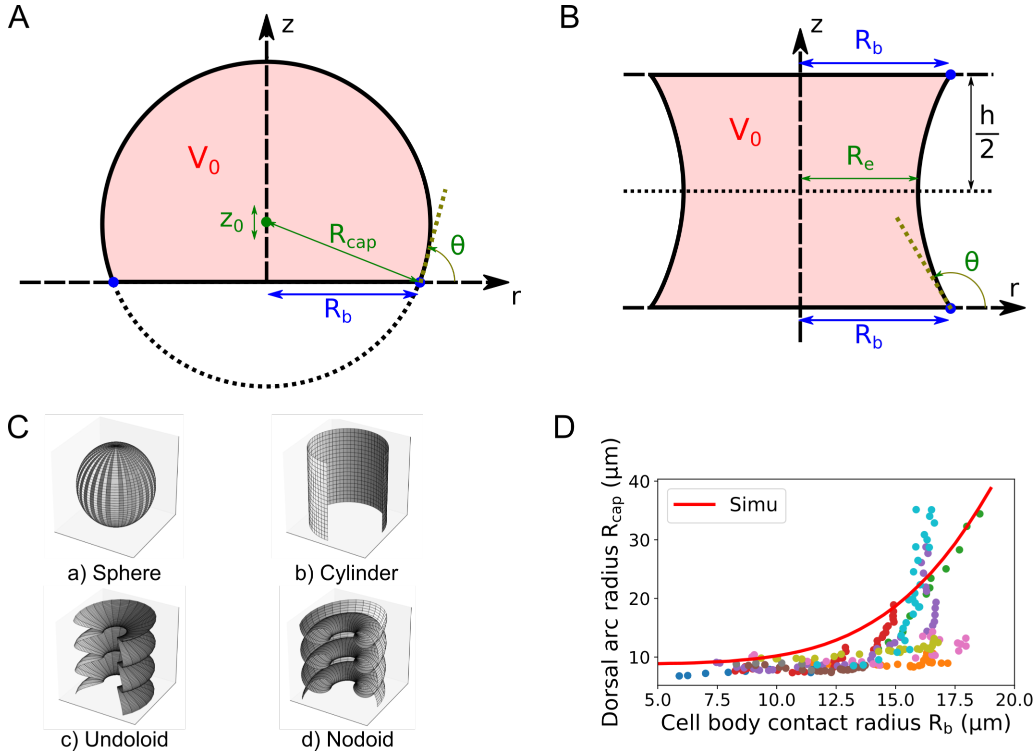


**Fig.S 5.** Geometrical description of cell spreading. A - Cell shape on a single plate is modeled as a spherical cap of main radius $R_{cap}$ and center $(0, z_0)$. These parameters are calculated so that the spherical cap volume is equal to $V_0$ and that the extreme contact point is at a given $r = R_b(t)$, which can be vary to predict cell body shape along spreading. B - Cell free edge between two plates is modelled by a constant mean curvature surface, symmetric around $z = 0$ and truncated by the plates at $z = \pm h/2$. C - Illustration of constant non-zero mean curvature (CMC) surfaces: (a) Sphere, (b) Cylinder, (c) Unduloid, and (d) Nodoid. D - Dorsal arc radius ($R_{cap}$ on Fig. A) as a function of cell body contact radius $R_b$ for the single plate geometry. Each color represent a single cell. Solid red line shows the result of simulations.

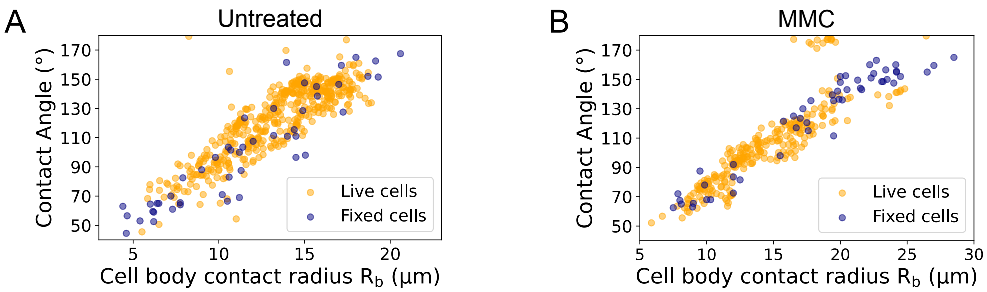


**Fig.S 6.** Comparison of three-dimensional cell shape in fixed (blue) and live (yellow) cells. A - Contact angle as a function of cell body contact radius for untreated cells. B - Contact angle as a function of cell body contact radius for mitomycin C-treated cells.

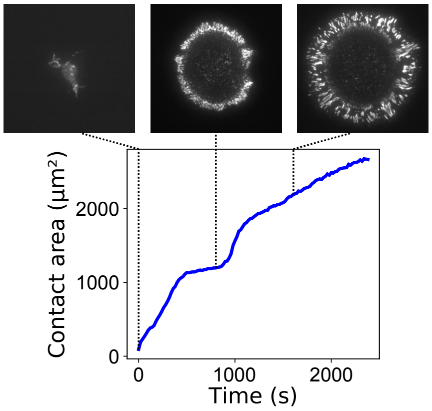


**Fig.S 7.** Rare event (n/N = 3/30) of cell spreading in two steps : spreading slow-down is followed by increase of spreading rate and second slow-down. Top row : TIRF imaging of paxillin-YFP along spreading. Left : Cell contact at t = 0s. Center : Cell contact at t=800s (first corona of paxillin clusters). Right : Cell contact at t=1600s (second corona of paxillin clusters). Bottom row : Cell contact area along spreading.

### Supplementary movies legend

**Movie 1.** Time-lapse of Ref-52 fibroblast expressing paxilin-YFP spreading on a bi-dimensional glass substrate, imaged via TIRF microscopy. Time is in min:sec. Scale bar: 5µm.

**Movie 2.** Time-lapse of Ref-52 fibroblast expressing paxilin-YFP spreading between two parallel glass plate. Bottom cell-substrate contact is imaged via TIRF microscopy. Time is in min:sec. Scale bar: 5µm.

**Movie 3.** Time-lapse of Ref-52 fibroblast expressing paxilin-YFP treated with mitomycin C spreading on a bi-dimensional glass substrate, imaged via TIRF microscopy. Time is in min:sec. Scale bar: 5µm.

**Movie 4.** Time-lapse of Ref-52 fibroblast spreading on a glass plate seen in profile, imaged via bright-field microscopy. Time is in min:sec. Scale bar: 5µm.

**Movie 5.** Time-lapse of Ref-52 fibroblast spreading between parallel glass plate seen in profile, imaged via bright-field microscopy. Time is in min:sec. Scale bar: 5µm.

**Movie 6.** Segmentation and markers of cell profile along spreading of a Ref-52 fibroblast spreading on a glass plate. Scale bar: 5µm.

**Movie 7.** Shape changes and markers of the profile of a spherical cap of constant volume, mimicking cell spreading on a bi-dimensional substrate.

**Movie 8.** Shape changes and markers of the profile of a constant mean curvature surface in contact with two parallel planes, mimicking cell spreading between parallel plates.

**Movie 9.** Three-dimensional representation of shape changes of a constant mean curvature surface in contact with two parallel planes, mimicking cell spreading between parallel plates.

**Movie 10.** Time-lapse of Ref-52 fibroblast expressing paxilin-YFP spreading on a bi-dimensional glass substrate, imaged via TIRF microscopy, showing two steps of spreading and two successive focal adhesions ring forming. Time is in min:sec. Scale bar: 5µm.
